# Roles and transcriptional regulation of the endogenous cellulases in association with substrate and cellular processes in a thermoacidophilic archaeon

**DOI:** 10.1101/2025.03.28.645846

**Authors:** Shuai Li, Qihong Huang, Yunfeng Yang, Pengju Wu, Jing Li, Yulong Shen, Jinfeng Ni

## Abstract

Microbes of the order *Sulfolobales* are potentially next generation platforms for cellulose degradation and utilization. However, the mechanisms of cellulose degradation and utilization remain unclear. In this study, we analyzed the enzymatic activity, localization, transcriptional regulation, and interplay of three endogenous cellulases Cel1, Cel2A, and LacS, and the association of transcription with cellular processes in the model thermophilic archaeon *Saccharolobus islandicus* REY15A. Overexpression strains based on a vector developed in this study as well as single and double deletion mutants of the three cellulase genes were constructed and analyzed. We reveal that Cel1 and Cel2A are membrane-associated, and a higher proportion of Cel2A is secreted into the medium than Cel1. The expression of all the three cellulases is induced by carboxymethylcellulose sodium (CMC). Furthermore, we found that the transcriptional levels of *cel1*, *cel2A*, and *lacS* are interdependent and *cel1* and *cel2A* levels apparently increased with *lacS* deletion and overexpression of an ABC transporter, suggesting of an intracellular oligosaccharide-dependent regulatory mechanism. We showed that a cell cycle transcription factor aCcr1 binds to the promoter of *cel1 in vitro*. Additionally, we accidentally found that the cellulase genes expression increases in the presence of uracil synthesis pathway and deletion of the cellulase genes facilitates cell growth in CMC-containing medium. This study reveals an ingenious and intricated mechanism for cellulose utilization to adapt to extreme environment in archaea and provide insights for engineering *Sulfolobales* archaea as biomass-degrading and utilization platforms.

## 1. Introduction

Lignocellulose is the most widely distributed carbohydrate in nature and represents an important renewable biomass resource with great application potentials (Ashokkumar et al., 2022; Reshmy et al., 2022). Efficient biomass utilization faces several technical challenges, particularly in the preprocessing and conversion processes. Common preprocessing methods include physical, chemical, and combined approaches. Although these conventional methods can facilitate biomass degradation to some extent, they are often associated with high energy consumption, stringent reaction conditions, and low conversion efficiency, which limit their widespread industrial application (Wu et al., 2021; Yan et al., 2020). To overcome these bottlenecks, increasing research efforts have focused on the role of microorganisms in biomass degradation (Malik & Javed, 2024; Mishra et al., 2017; Sperandio & Filho, 2021; Woo et al., 2014; Ying et al., 2024). Meanwhile, acid-thermal pretreatment, as an effective method, can disrupt the crystalline structure and physical barriers of lignocellulose (Guo et al., 2022; Zhang et al., 2023), typically requiring subsequent cooling and neutralization, etc.

The microbes of *Sulfolobales*, which thrive in extreme acidic and thermal environments could offer significant advantages in biomass biodegradation and fermentation. On one hand, biomass degradation and fermentation can be combined (one-pot synthesis) by which alkaline neutralization and cooling processes after biomass pretreatment can be eliminated, greatly reducing cost. On the other hand, these archaea can simultaneously utilize both pentose and hexose sugars. In addition, the thermophilic and acidophilic growing conditions of *Sulfolobales* archaea can avoid contamination in industrial fermentation (Crosby et al., 2019; Pfeifer et al., 2021; Schocke et al., 2019). In recent years, advances in genetic manipulation and understanding of archaea biology have facilitated metabolic pathway modifications through genome editing techniques (Müller, 2014). However, the cellulolytic mechanisms of these microbes remain mysterious (Cowan et al., 2024; De Lise et al., 2023; Lewis et al., 2021).

*Saccharolobus islandicus* REY15A as a model archaeon of *Sulfolobales* has a well-developed genetic manipulation system, including the establishment of auxotrophic selection markers, inducible overexpression plasmids, and gene editing tools based on the endogenous CRISPR-Cas system (Deng et al., 2009; Guo et al., 2011; Li et al., 2016; Peng et al., 2012; Zheng et al., 2012). In *Sa. solfataricus* (formerly *Sulfolobus solfataricus*), a close relative of *Sa*. *islandicus*, four cellulases SsoCelS (*sso2534*), SSO1354 (*sso1354*), SSO1949 (*sso1949*), SsoLacS (*sso3019*) were predicted (She et al., 2001). The colony of *Sa. solfataricus* on carboxymethylcellulose sodium (CMC)-containing plates formed a hydrolysis zone and *sso2534* transcript was highly induced by CMC, demonstrating that SsoCelS may have endoglucanase activity *in vivo* (Limauro et al., 2001). Recombinant SSO1949 protein exhibited endoglucanase activity *in vitro* (Huang et al., 2005). It was suggested that either SSO1354 or SSO1949 originated from a duplication/insertion event mediated by insertion sequences (ISs) (Girfoglio et al., 2012). SSO1949 has an optimal pH of 1.8 and an optimal temperature of 80°C (Huang et al., 2005), while SSO1354 undergoes glycosylation, with an optimal pH of 3.5 and an optimal temperature of 95°C (Maurelli et al., 2008). Additionally, it is likely that SSO1354 functions by anchoring to the cell membrane (Girfoglio et al., 2012; Maurelli et al., 2008). SsoLacS, a reporter gene often used in genetic research, exhibits β-galactosidase activity and can hydrolyze β-D-gluco-, fuco-, and galacto-sides. It is also capable of degrading G2-G5 oligosaccharides. SsoLacS can synergistically interact with ABC transporter substrate binding protein SSO3053 (which can bind cellodextrin from G2 to G8), forming an efficient cellodextrin degradation system (Lalithambika et al., 2012; Nucci et al., 1993; Wang et al., 2023; Zheng et al., 2012). Nevertheless, it is still obscure about how the cellulases and other proteins work in cellulose degradation *in vivo*.

Genomic annotation identified three predicted cellulases in *Sa. Islandicus* REY15A, Cel1, Cel2A, and LacS, homologs to SsoCelS (*sso2534*), SSO1354 (*sso1354*)/SSO1949 (*sso1949*), and SsoLacS (*sso3019*), respectively (Guo et al., 2011). In this study, we analyzed the enzymatic activity, cellular localization, and interplay of the three cellulases. The transcriptional regulation in response to carboxymethyl cellulose (CMC), growth and nutrient conditions was also investigated. A novel constitutive expression vector pSeAP was constructed. We revealed that Cel1 and Cel2A are membrane-associated proteins, with Cel2A exhibiting a higher secretion ratio than Cel1. Additionally, aCcr1, a cell cycle repressor identified in our previous study (Yang et al., 2023) was found to bind *cel1* promoter. The transcription of *Sa. islandicus* cellulases is upregulated in the presence of CMC. Furthermore, we found that the transcriptional regulation of these cellulases is probably influenced by intracellular oligosaccharide levels.

## 2. Materials and methods

### 2.1 Strains and growth conditions

*Sa. islandicus* REY15A was grown aerobically at 75℃ in GTV medium containing mineral salt, 0.2% (w/v) glucose (G), 0.2% (w/v) tryptone (T), and a mixed vitamin solution (V). E233 was obtained by knocking out the *pyrEF* gene in *Sa. islandicus* ReY15A. The medium was adjusted to pH 3.3 with sulfuric acid, as described previously (Deng et al., 2009; Yang et al., 2023). E233 and other strains without plasmids were grown in medium supplemented with 0.01% (w/v) uracil (U). CMC-TV (U) medium containing 0.2% (w/v) CMC (Macklin, Shanghai, China) was used for cellulase induction and growth curve measurement. The other carbon sources used for cultivation are D-glucose, D-cellobiose and Microcrystalline Cellulose (MMC) (Sinopharm, Shanghai, China). CMC plates were prepared using gelrite (0.8% [w/v]) by mixing 2 × CMC-TV and an equal volume of 1.6% gelrite. The strains constructed and used in this study are listed in the Supplementary Table S1.

### 2.2 Construction of constitutive expression vector pSeAP

DNA fragment of the Alba promoter containing *Sph*I and *Nhe*I cleavage sites were obtained by PCR using genomic DNA of *Sa. islandicus* E233 and Primers 1-2 (Supplementary Table S2). The PCR fragment and pSeSD DNA was both digested with *Sph*Ⅰ and *Nhe*Ⅰ, then the vector and DNA fragments were ligated by T4 ligase, yielding the expression vector pSeAP. For construction of recombinant expression vectors (Supplementary Table S1, Nos.19-23), the plasmids were digested with *Nhe*Ⅰ and *Sal*Ⅰ for pSeAP, and *Nde*I and *Sal*I for pSeSD, respectively, then the target gene was ligated to the above linearized vector using seamless cloning method (Xia et al., 2019). The primers used for constructing the overexpression vectors of Cel1, Cel2A, LacS, and SBP are listed in Supplementary Tables S2 (Primers 3-12).

### 2.3 Protein structure prediction by Alphafold 3

The protein sequences of Cel1, Cel2A, and LacS of *Sa. islandicus* REY15A were downloaded from NCBI website (https://www.ncbi.nlm.nih.gov/), the sequences were uploaded to Alphafold 3 online website (https://golgi.sandbox.google.com/about) for prediction.

### 2.4 Preparation of membrane fraction by ultracentrifugation

Cells grown in GTV medium (OD_600_ of 0.4∼0.6) were harvested and the medium supernatant was concentrated by centrifugation with a 30 kDa cutoff membrane. The pellet was suspended in 50 mM Tris-HCl (pH 8.0) and subjected to ultrasonication on ice. The sample was centrifugated at 2,000×g for 10 min, then the supernatants was centrifuged at 240,000×g for 20 min (Beckman Coulter Life Sciences). After centrifugation, the supernatant was collected as soluble cytoplasma fraction. The precipitate was resuspended with the same buffer as membrane fraction.

### 2.5 Gene knock out

Construction of the knockout strains was carried out according to the endogenous CRISPR-Cas-based genome editing method (Li et al., 2016). The 40 nt DNA sequence downstream of the CCN or TCN motif was selected as the protospacer, and annealing of the two complementary oligonucleotides yielded a DNA fragment containing the designed spacer. The spacer fragment and the digested vector were ligated by DNA ligase to yield the mini-CRISPR plasmid. The donor DNA fragment was generated by splicing and overlapping extension (SOE)-PCR with *ApexHF* HS DNA Polymerase FS (Accurate Biotechnology, Hunan, China), following the reported procedure (Horton et al., 1989). Then the donor DNA fragment was digested with *Sph*I and *Xho*I and inserted into the mini-CRISPR plasmid digested with same restriction enzymes to obtain the genome editing plasmids (Supplementary Table S1, No. 24-27). The sequences of all constructed plasmids were verified by DNA sequencing (Sangon Biotech, Shanghai, China). The cells of *Sa. islandicus* were transformed with the plasmid by electroporation (Schleper et al., 1992). The in-frame deletion strains were screened and verified by PCR and subsequent sequencing of the PCR products. The gene editing plasmid was removed by *pyrEF* counter selection in the presence of uracil and 5-fluoroorotic acid (5-FOA). The primers used for constructing the knockout plasmids of Cel1, Cel2A, LacS, and SBP are listed in Supplementary Tables S2 (Primers 13-36).

### 2.6 Growth curve measurement and CMC induction assay

Cells were cultured in GTV or GTVU for revival, allowing the cells to grow to the logarithmic growth phase prior to further analysis. The cells were inoculated into 40 ml medium to a final estimated OD_600_ of 0.04 and the growth was monitored using spectrometer (Persee, Beijing, China). For CMC induction assay, cells at logarithmic growth phase were harvested at 0.5 OD (approximately 2.5 × 10^8^ cells), the samples were centrifuged at 7,000 × *g* for 1 min at room temperature. The supernatant was discarded and the cell precipitate was resuspended using 5 ml of CMC-TV or CMC-TVU medium, and then the cells were continuously cultured at 75°C with shaking for 12 h.

### 2.7 Quantitative reverse transcription PCR (RT-qPCR)

Cells were centrifuged at 6,000×*g* for 1 min and the SparkZol Reagent (SparkJade, Shandong, China) was added. Total RNA was extracted according to the manufacturer’s instructions. The first-strand cDNA was synthesized by reverse transcription of RNA using *Evo M-MLV* RT Mix Kit (Accurate Biotechnology, Hunan, China) according to the manufacturer’s protocol. The mRNA levels were evaluated by RT-qRCR using the SYBR Green Premix *Pro Taq* HS qPCR Kit (Accurate Biotechnology, Hunan, China) on the CFX96 Touch™ Real-Time PCR System (Bio-Rad, Hercules, CA, USA). The two-step PCR amplification standard procedure was as follows: 95℃ for 30 s, followed by 40 cycles of PCR that consisted of 95℃ for 5 s and 60℃ for 30 s. The relative transcription level of the target gene was calculated using the comparative threshold cycle (CT) method 2^(-ΔΔCt) (Schmittgen & Livak, 2008), and the *tbp* was used as reference. The primers used in this study are listed in Supplementary Tables S2 (Primers 39-46).

### 2.8 β-Galactosidase assay

The β-galactosidase assay of the reporter gene *lacS* (β-glycosidase) was basically performed using ONPG (o-Nitrophenyl-β-D-galactopyranoside) method (Peng et al., 2009). The cells in the logarithmic growth phase were collected and re-suspended in 50 mM Tris-HCl (pH 8.0). The cell suspensions were sonicated to yield crude cellular extracts. After removing cell debris by centrifugation (13,000 rpm for 30 min at 4℃), the cell lysates were suitable for ONPG assay. Protein contents of the cellular extracts were determined using Bradford Protein Assay Kit (Beyotime, Shanghai, China). Samples were either assayed immediately or stored at −80℃. For each assay, 10–50 μL supernatant was transferred to a microcentrifuge tube containing 450–490 μL reaction buffer with 2.8 mM ONPG and 50 mM sodium phosphate, pH 6.5, in a total of 500 μL. Samples were incubated at 75℃ for 10 min and the reaction was stopped by adding an equal volume of 1 M sodium carbonate. Yields of o-nitrophenol were determined by measuring at 420 nm using visible spectrophotometer (Persee, Beijing, China). One specific unit was defined as 1 mmol o-nitrophenol produced per min per mg total protein.

### 2.9 Western-blotting

Antibody against TBP (TATA-box Binding Protein) was prepared utilizing synthetic specific peptides (amino acids 18–31, SIPNIEYDPDQFPG for TBP). TBP antibody was produced in rabbits (HuaAn Biotechnology, Hangzhou, China). After transformed, cells grown to the logarithmic phase were harvested and the pellet was re-suspended in 50 mM Tris-HCl (pH 8.0) and subjected to ultrasonication on ice. The supernatant was concentrated by centrifugation with a 30 kDa cutoff membrane. The resulting total cellular protein or supernatant fraction were supplemented with 5 × loading buffer and boiled for 10 min. Proteins were fractionated by 15% sodium dodecyl sulphate-polyacrylamide gel electrophoresis (SDS-PAGE) and transferred onto polyvinylidene difluoride (PVDF) membranes at 30 mA for 16 h at 4℃. The membranes were blocked by incubating in 5% skim milk for 2 h at room temperature. When incubating multiple antibodies, the membranes were cut and incubated, with primary anti-His or anti-TBP antibody separately and subsequent secondary goat anti-rabbit HRP conjugate antibody (TransGen Biotech, Beijing, China) according to the manufacturer’s instructions. The signals were visualized using an Amersham ImageQuant 800 (Cytiva, USA).

### 2.10 Native-PAGE and CMC degradation activity assay

Proteins were separated by Native-PAGE in gels containing 0.1% (w/v) CMC. The electrophoresis was run at 200 V for 90 min, subsequently the gel was first rinsed with PBS buffer (pH 7.4) for 10 min at room temperature, followed by replacing it with a new PBS buffer (pH 7.4) and incubating for 30 min at 75℃. The gel was treated with citric acid-sodium citrate buffer (pH 3.4) and incubated for 30 min at 75℃. The gel was stained with 0.1% (w/v) Congo red solution for 30 min and washed with 1 M NaCl for 10 min. For revealing cellulase activity, basically it was done as described (Ni et al., 2010) with modification. The overexpression or knockout strains were grown for 2 days at 75℃ on plates containing 0.2% (w/v) CMC. Then the plates were washed with 75% ethanol, stained with 0.1% (w/v) Congo red solution for 30 min and finally washed with 1 M NaCl for 10 min. Cellulase activity (CMCase) was visualized as a hydrolysis zone around colony against the red background.

### 2.11 Electrophoretic mobility shift assay (EMSA)

Substrates used in EMSA experiments were generated by annealing the complementary oligonucleotides with 5’ FAM-labelled oligonucleotides (Supplementary Table S2, Primers 47-52). The reaction mixture (20 μL) containing 2 nM of the FAM-labelled substrates and different concentrations of aCcr1 proteins was incubated at 37℃ for 30 min in binding buffer (25 mM Tris–HCl, pH 8.0, 25 mM NaCl, 5 mM MgCl_2_, 10% glycerol, 1 mM dithiothreitol). After the reaction, samples were loaded onto a 10% native PAGE gel buffered with 0.5 × Tris–Borate–EDTA (TBE) solution. DNA–protein complexes were separated at 200 V for 60 min. The resulting fluorescence was visualized by an Amersham ImageQuant 800 biomolecular imager (Cytiva).

## 3. Results

### 3.1 Construction of a constitutive expression vector pSeAP

In the model archaeon *Sa. islandicus*, the arabinose inducible vector pSeSD is commonly used for efficient protein expression (Peng et al., 2012). To avoid the effect of different sugars on protein expression, we aimed to construct a constitutive expression vector. We screened the transcriptomic data of *Sa. islandicus* REY15A for high constitutive expression promoter and identified the promoter (*P_alba_*) of the chromosomal DNA-binding protein SisAlba (SiRe_1125) as candidate (Guo et al., 2011; Yang et al., 2023). *P_alba_* is 77 bp in length and contains predicted BRE, TATA box, BoxA, and Shine-Dalgarno (SD) sequences (Condò et al., 1999; Peng et al., 2012; Tolstrup et al., 2000; Torarinsson et al., 2005) (Figure 1A). The *P_araS_* in pSeSD was replaced with *P_alba_*, yielding a constitutive overexpression vector pSeAP (Figure 1B), which contains a multiple cloning sites (MCS) and a sequence for a His-tag at the downstream of MCS (Figure 1C). To evaluate the transcriptional and protein expression efficiency using pSeAP, we constructed two recombinant strains carrying pSeSD-LacS and pSeAP-LacS, respectively. RT-qPCR and β-Galactosidase assay were used to assess the differences in transcription and enzymatic activity between the two strains. As expected, the transcription of *lacS* in inducible expression strain was low under non-inducing conditions (D-glucose), but significantly upregulated under D-arabinose induction (Figure 1D). In contrast, pSeAP-LacS enabled *lacS* constitutively expressed under both D-glucose and D-arabinose conditions, with transcription levels reaching 135.43% and 142.10%, respectively, of that in the strain carrying pSeSD-LacS with adding D-arabinose induction (Figure 1D). β-Galactosidase activity assay showed that strain harboring the inducible vector has very low activity with D-glucose but high activity with D-arabinose (19.53-fold), while the strain carrying the constitutive vector exhibited significantly higher galactosidase activity in both media, reaching 164.50% and 158.90%, respectively, of that in the inducible strain under D-arabinose induction (Figure 1E). Overall, the pSeAP vector enables efficient transcription and high expression under non-inducing conditions, making it a valuable tool for constitutive gene expression in *Sa. islandicus*.

**Fig. 1.**
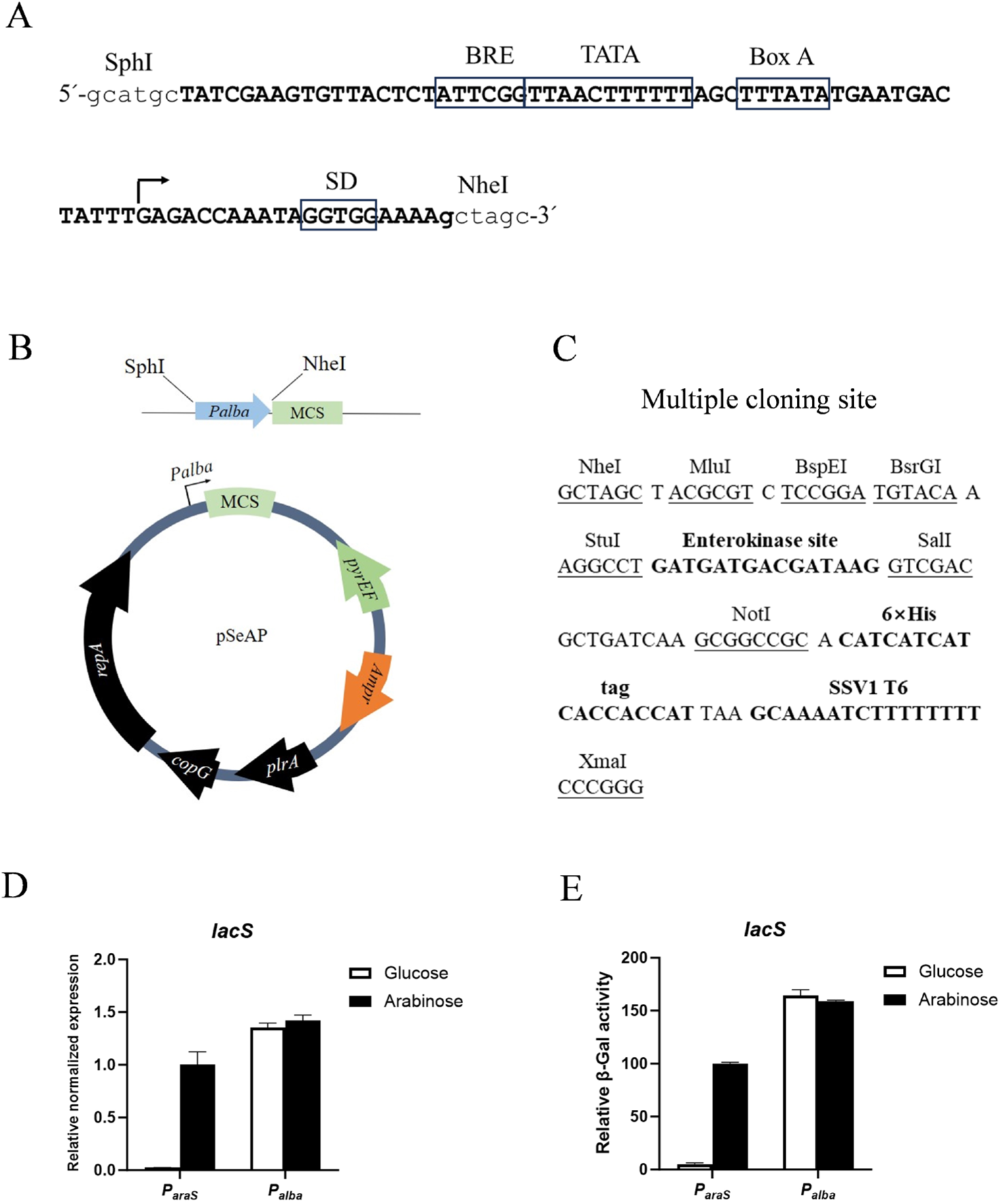
Construction and verification of a constitutively overexpression vector pSeAP for *Sa*. *islandicus.* (**A**) Sequence features of the *P_alba_* promoter. Capital letters show the sequence of the promoter, with putative promoter elements boxed and indicated above the sequence motifs. The lowercase letters are the restriction sites flanking the promoter. BRE, transcription initiation factor B recognition element; TATA, AT-rich sequence motif in promoters serving as the binding site for TATA box-binding protein TBP; Box A, signal sequence which is essential for transcription efficiency; The arrow indicated the transcription start site; SD, ribosome-binding sequence Shine-Dalgarno. (**B**) Schematics maps of the constitutive expression vector pSeAP. (**C**) The multiple cloning site sequence of pSeAP. (**D**) Comparison of the transcript levels of *lacS* between two recombinant strains carrying inducible pSeSD-LacS and constitutive pSeAP-LacS. Cells were cultured in GTV (glu) or ATV (ara) and collected at an OD_600_ of 0.2 for RNA extraction and RT-qPCR. The RT-qPCR was performed using TBP gene as the internal reference. (**E**) Comparison of the β-galactosidase activities of the two. Cells were cultured and were harvested as in (**D**). All assays were conducted in triplicates.

### 3.2 Structural prediction and expression of endogenous cellulases in Sa. Islandicus

We named the cellulase homologs SiRe_0332, SiRe_0791, and SiRe_2224 in *Sa. islandicus* (Guo et al., 2011) as SisCel1, SisCel2A, and SisLacS, respectively. Firstly, we predicted their structures by AlphaFold 3. As shown in Figure 2A, both Cel1 and Cel2A have a N-terminal long helix domain, while LacS does not. Next, we cloned *cel1*, *cel2A*, and *lacS* into pSeAP generating vectors pSeAP-Cel1, pSeAP-Cel2A, and pSeAP-LacS. After transformation into *Sa. islandicus* E233, we obtained strains carrying each of the expression vectors. The strains were cultivated in GTV medium and the cells were collected for analysis (Figure 2B and 2C). Western blotting was used to detect the presence and distribution of these proteins. The theoretical molecular weights of Cel1, Cel2A, and LacS are 38.2 kDa, 38.9 kDa, and 58.2 kDa, respectively. In the total cellular samples, both Cel1 and Cel2A appeared at approximately 45.0 kDa and 57.0 kDa, respectively, larger than their theoretical molecular sizes (Figure 2B). While LacS appeared as a band that was consistent with its theoretical molecular size (Figure 2B). In the medium supernatant, both Cel1 and Cel2A were detected, but LacS was not detected (Figure 2B). The molecular size of Cel1 was identical to that observed in the total cellular protein, whereas Cel2A appeared as two distinct bands, one matching the size observed in the total cellular sample and one close to its theoretical size. Notably, although the same expression vector of pSeAP was used, the LacS showed higher expression levels compared to Cel1 and Cel2A, as evidenced by the internal reference protein TBP (Figure 2B). Then activity staining was performed using Native-PAGE gels containing CMC. Surprisingly, the Cel1 expression strain did not show active band in either the total cellular protein or the supernatant (Figure 2C). In contrast, CMC-degrading activities were observed both in total cellular protein and supernatant of the Cel2A expression strain, and an apparent activity band predominantly presented in the culture supernatant. As expected, no CMC-degrading activity was detected for the LacS expression strain (Figure 2C), which is in agreement with that LacS as a β-glucosidase was unable to hydrolyze CMC.

**Fig. 2.**
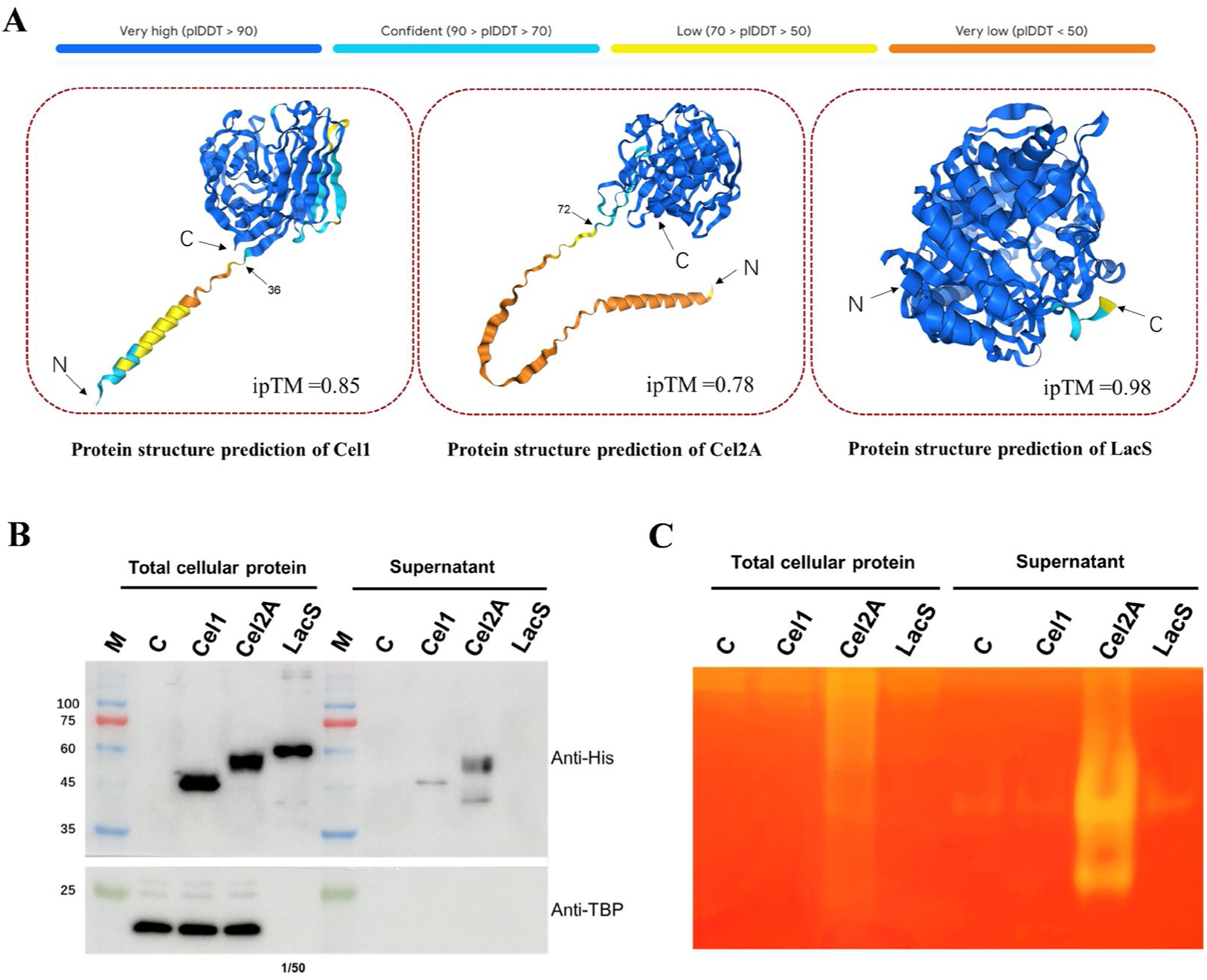
Structural prediction, ectopic expression, and activity analysis of the endogenous cellulases from *Sa*. *islandicus.* (**A**) Structures of Cel1, Cel2A, and LacS. The structures were predicted by Alphafold 3. The above shows prediction confidence schematic. IpTM (Interface predicted template modeling) indicates the accuracy of the prediction. The maximum value is 1, and the higher the value, the higher the confidence of the predicted structure. (**B**) The protein levels of Cel1, Cel2A and LacS in *Sa*. *islandicus*. Cells of E233 were transformed with pSeAP, pSeAP-Cel1, pSeAP-Cel2A, and pSeAP-LacS, respectively. After grown on GTV medium to OD_600_ of 0.4∼0.6, cells were harvested and the pellet was suspended in 50 mM Tris-HCl (pH 8.0) and subjected to ultrasonication on ice. The medium supernatant was concentrated by centrifugation in a concentrator tube with 30 kDa cutoff membrane. Cells for LacS expression was sampled as for Cel1 and Cel2A but for the analysis the sample was diluted by 50-fold, thus the TBP signal in the LacS sample well is weaker and invisible. M, Protein standard marker; C, control of cell harboring the empty plasmid pSeAP. Total cellular protein and proteins in the concentrated medium were analyzed by SDS-PAGE and subsequent Western-blotting against anti-His and TBP antibodies, respectively. TBP, TATA-box binding protein. (**C**) Enzymatic activity analysis of samples of Cel1, Cel2A, and LacS overexpression strains by Native-PAGE and zymography. Total cellular extracts and concentrated medium fractions were subjected to Native-PAGE on a gel containing 0.1% CMC and the cellulase activity was revealed by zymography by staining with Congo red.

To further investigate the role of these cellulases in *vivo*, we constructed and obtained the individual knockout stains of *cel1* (Δ*cel1*), and *cel2A* (Δ*cel2A*). The deletion strain of *lacS* (Δ*lacS*) is available in our lab stock (E233S). Three knockout stains and three cellulase expression strains were cultured on CMC-containing plates. As shown in Figure 3A, Δ*cel1* exhibited a hydrolysis zone similar to that of E233, the parent strain, whereas Δ*cel2A* showed no detectable hydrolysis zone. As expected, Δ*lacS* displayed hydrolysis zone size with no significant difference with that of E233. For the overexpression strains (Figure 3B), strains carrying pSeAP-Cel1 and pSeAP-LacS did not produce hydrolysis zone distinct from that of strain carrying pSeAP. However, E233/pSeAP-Cel2A showed a markedly larger zone compared to strain carrying pSeAP. These results suggest that *cel2A* is the only active endoglucanase present in *Sa. islandicus* and indicate that Cel2A plays a critical role in cellulose degradation.

**Fig. 3.**
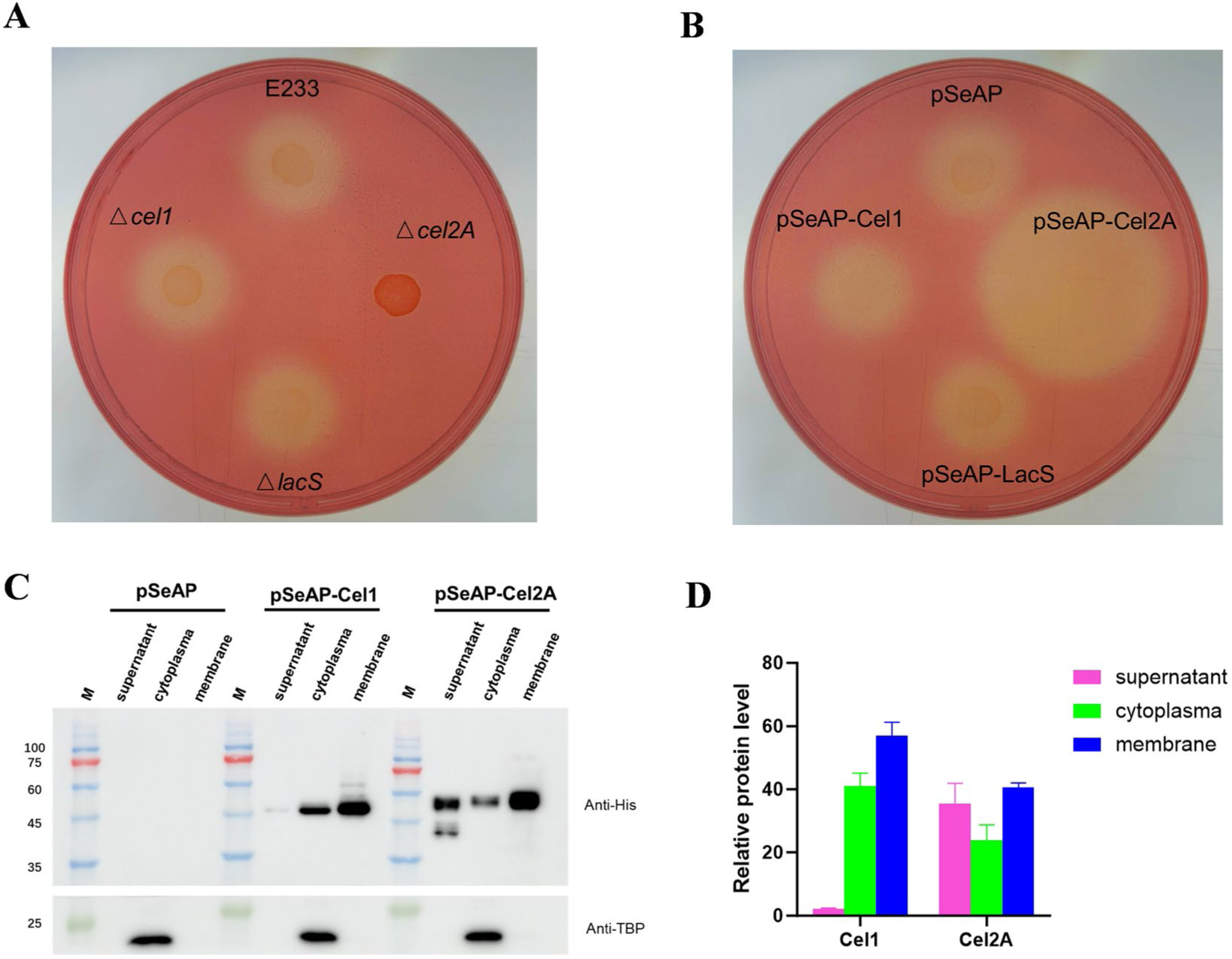
Congo red plate assay and subcellular localization of the three cellulases. (**A**) Activity assay of the parent strain E233 and the knockout strains by Congo red staining on plates containing 0.2% CMC. (**B**) Activity assay of control and the overexpressing strains of *cel1*, *cel2A*, and *lacS* by Congo red staining. (**C**) Localization of Cel1 and Cel2A in cells of the overexpression strains. The culture samples were fractioned into medium supernatant, cytoplasma, and membrane fractions. The proteins were detected by SDS-PAGE and subsequent Western-blotting against anti-His antibody and anti-TBP antibodies, respectively. The medium supernatant was concentrated by centrifugation in a concentrator tube with 30 kDa cutoff membrane. After collecting the cell samples by centrifugation, they were first sonicated on ice and then separated into cytoplasma and membrane fractions through ultracentrifugation. TBP, TATA-box binding protein. M, Protein standard markers. A representative gel image was shown. (**D**) Quantification of the results in (C) using ImageJ. All assays were conducted in triplicates.

To determine subcellular localization of Cel1 and Cel2A, we fractionated the samples of the control (pSeAP) and two expression strain cells (pSeAP-Cel1 and pSeAP-Cel2A) into different parts using low-speed and ultracentrifugation. Western blotting analysis combined with ImageJ-based quantification revealed that both Cel1 and Cel2A were predominantly membrane-associated, with 56.94% and 40.54% of their total amounts, respectively (Figure 3C-D). Cel2A had a significantly higher ratio in the supernatant, accounting for approximately 35.56%, whereas the supernatant of Cel1 was low, accounting for only 2.11% of its total protein. In cytoplasma fraction, Cel1 and Cel2A accounted for 40.94% and 23.91% of their total proteins, respectively (Figure 3C-3D). These results indicate that both Cel1 and Cel2A are primarily membrane-bound proteins and Cel2A is also secreted or released to the medium at a higher proportion.

### 3.3 Transcription of the cellulase genes is regulated by the cell cycle regulator aCcr1

Transcription factors play a crucial role in the regulation of cellulase gene expression, such as in *Trichoderma reesei* and *Talaromyces pinophilus* (Aro et al., 2001; Cziferszky et al., 2002; Furukawa et al., 2009; Liao et al., 2018; Saloheimo et al., 2000). In 2023, our research group reported a RHH-family transcription factor aCcr1 in *Sa. islandicus*. Overexpression of aCcr1 enlarged cells and reduced growth rate. Interestingly, *cel1* is among the highly downregulated genes upon aCcr1 overexpression (Yang et al., 2023), implying that the transcription of *cel1* in *Sa. islandisus* is associated with cell cycle or cell growth. To verify if the cellulase genes could be regulated by aCcr1, we performed EMSA to assess binding ability of aCcr1 to the promoters. The results showed that aCcr1 specifically binds *P_cel1_* but not *P_cel2A_* or *P_lacS_* (Figure 4A and B).

**Fig. 4.**
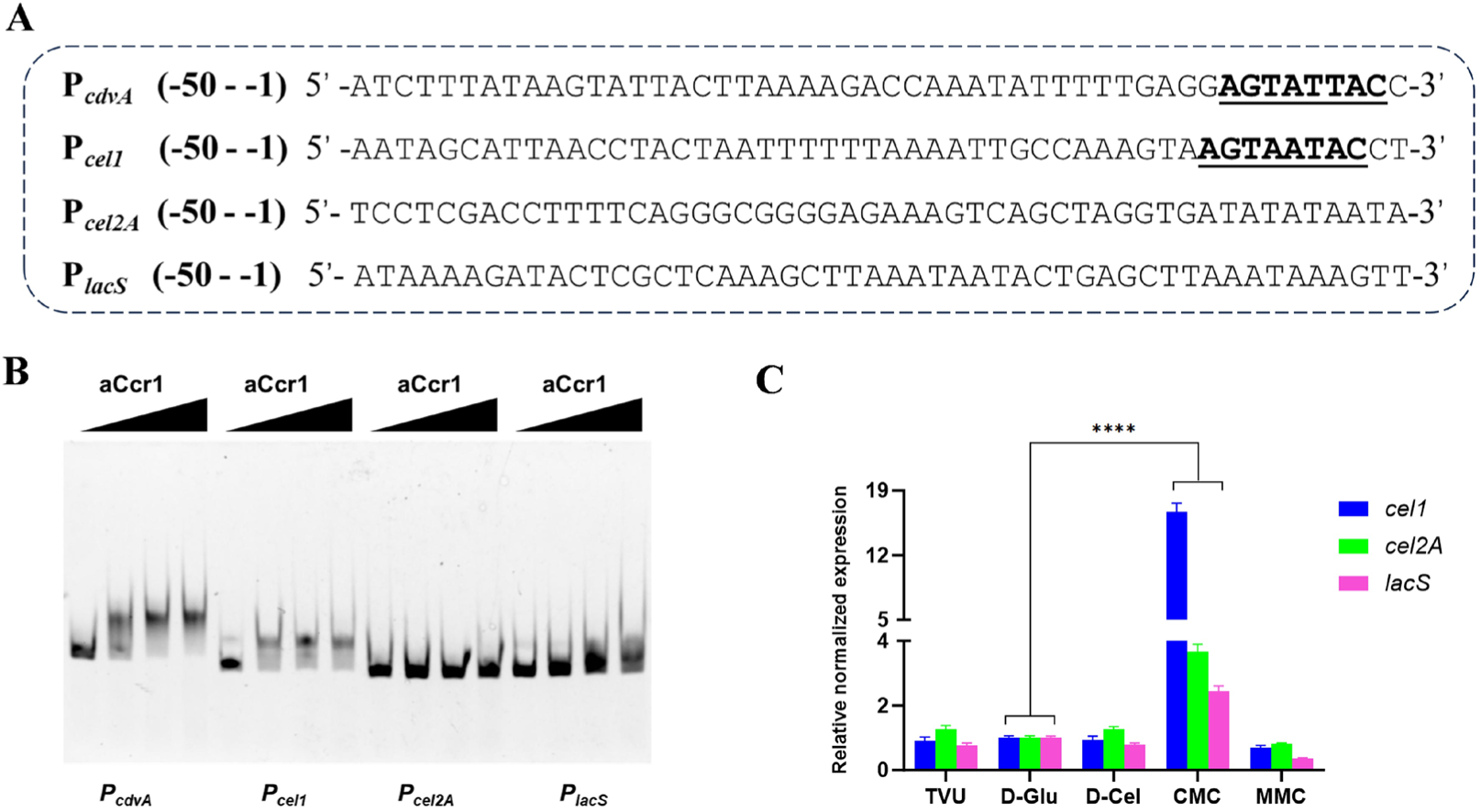
Transcriptional regulation of cellulases by transcription factor aCcr1 and induction by carbon sources in *Sa*. *islandicus.* (**A**) Promoter sequence and putative aCcr1 binding motif of *cdvA*, *cel1*, *cel2A*, and *lacS*. Bold text and underlined indicates predicted motifs potentially bound by aCcr1. (**B**) Assay of aCcr1 binding to the promoters of *cdvA*, *cel1*, *cel2A*, and *lacS*. P*cdvA* was used a positive control. DNA substrates were made with oligonucleotides labelled with FAM at the 5’-ends and the corresponding complementary oligonucleotides for EMSA. The reactions were performed at 37℃ for 30 min and the samples were analyzed on a 10% native-PAGE gel (see ‘Materials and Methods’). The reaction mixture contained 2 nM of FAM-labelled substrate and 0, 0.25, 0.5, and 1.0 μM aCcr1. Representative image is shown. (**C**) Transcription of cellulases of *Sa*. *islandicus* E233 in the presence of different carbon sources. TVU, basal medium with no carbon source; D-Glu, D-Cel, CMC, and MMC, TVU medium supplement with D-glucose, D-cellobiose, CMC, and MMC, respectively. Cells were harvested when grown to logarithmic phase and subjected to RNA extraction. The RT-qPCR experiments were performed in triplicates, with TBP gene as the internal reference.

### 3.4 Transcription of the cellulase genes is induced by CMC

In addition, cellulase gene expression is also induced by cellulose in other microorganism (Ilmén et al., 1997; Li et al., 2022; Yan et al., 2021; Zhou et al., 2012). For example, in *Myceliophthora thermophila*, Avicel can effectively induce the production of the cellulases (Li et al., 2022). To explore the effect of carbon sources on the transcription of cellulase genes in *Sa. islandicus*, we cultured the strain in media containing different carbon sources and analyzed the transcriptional levels of the three cellulase genes using RT-qPCR. Compared to D-glucose, all three cellulase genes were upregulated when CMC was used as carbon source. Among them, *cel1* was the highest upregulated, with a 15.66-fold increase compared to D-glucose. *cel2A* was upregulated by 2.68-fold, while *lacS* showed a 1.44-fold increase (Figure 4C). In contrast, no significant upregulation of the three cellulase genes was observed in the carbon-free control group (TVU) or in cultures supplemented with cellobiose and microcrystalline cellulose (MMC). Overall, our findings indicate that CMC can simultaneously induce the transcriptional upregulation of *cel1*, *cel2A*, and *lacS*. The mechanism about how CMC induces the transcription needs further investigation.

### 3.5 The transcription of cellulase genes is possibly regulated by intracellular oligosaccharides

In cellulase-producing microorganisms, the transcription of cellulase system genes can be regulated by multiple mechanisms, including interplay among different cellulases and intracellular sugar level (Chen et al., 2013; Fowler & Brown, 1992; Li et al., 2022; Zhou et al., 2012). To understand whether this regulation exists in *Sa. islandicus*, RT-qPCR was performed to analyze the change in transcriptional level of other cellulase genes in each of the deletion strains. The results showed that in Δ*cel1*, *cel2A*, and *lacS* transcription were downregulated to 36.41% and 16.93% of that in E233, respectively. In Δ*cel2A*, *cel1* and *lacS* transcription levels were reduced to 22.66% and 21.46% of that in E233, respectively. Conversely, in Δ*lacS*, *cel1*, and *cel2A* were upregulated to 202.84% and 169.44% of that in E233, respectively (Figure 5A). This pattern is similar to the phenomenon observed in *Penicillium decumbens*, where the suppression of intracellular β-glucosidase transcription enhances extracellular cellulase production. In *P. decumbens*, the transcriptional change is caused by accumulation of intracellular cellobiose due to reduced intracellular β-glucosidase activity (Chen et al., 2013). We hypothesize that the accumulation of LacS substrates might induce the upregulation of *cel1* and *cel2A*. LacS was reported to have hydrolytic activity on cello-oligosaccharides ranging from G2 to G5 (Lalithambika et al., 2012; Nucci et al., 1993). We speculate that *lacS* deletion leads to the accumulation of intracellular cellooligosaccharides, thereby inducing the transcriptional upregulation of *cel1* and *cel2A*.

**Fig. 5.**
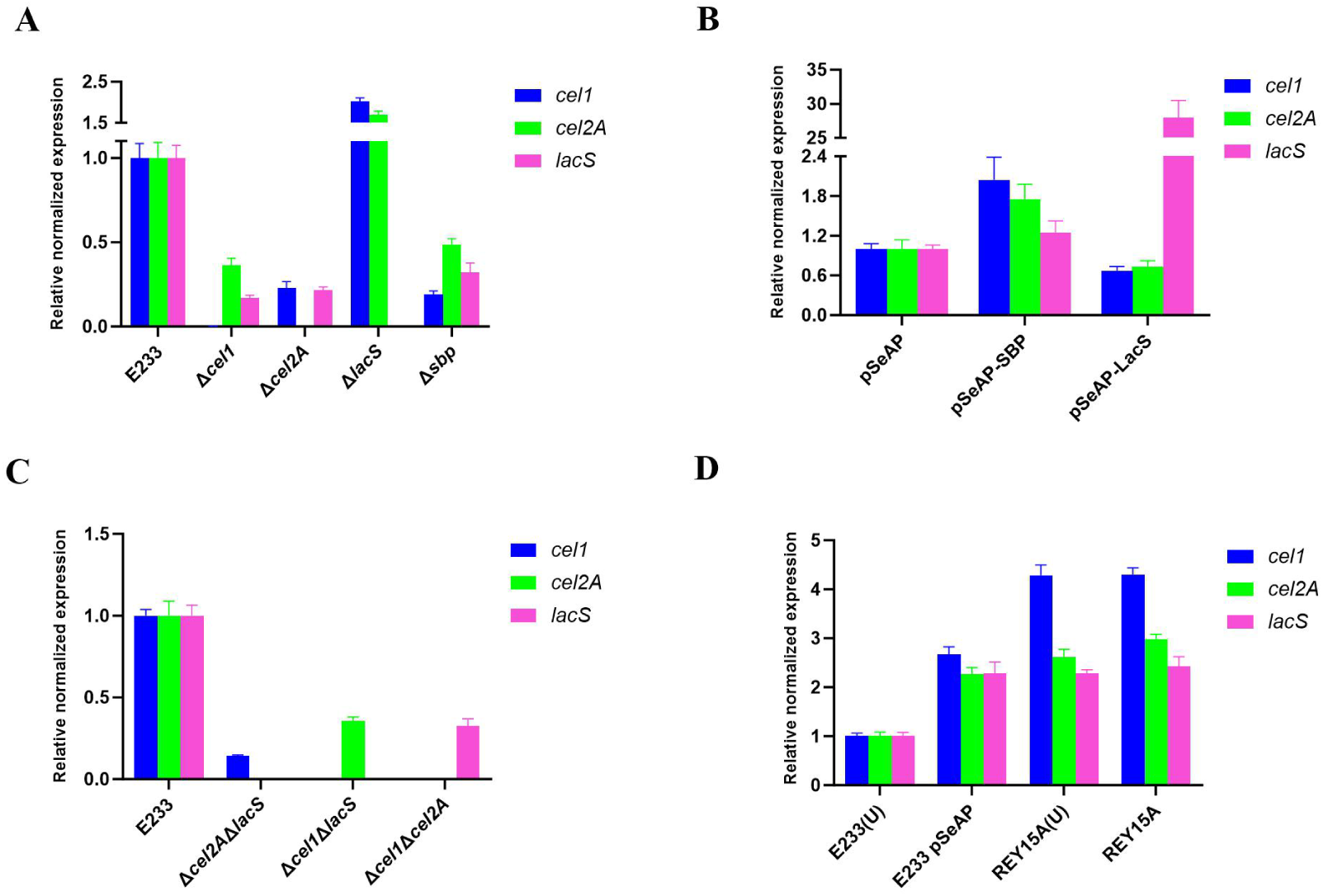
Changes of the relative transcription levels of *cel1*, *cel2A*, and *lacS* in CMC medium analyzed by RT-qPCR. (**A**) Single knockout strains of E233Δ*cel1*, E233Δ*cel2A*, E233Δ*lacS* and E233Δ*sbp*. (**B**) Overexpression strains of E233/pSeAP-LacS and E233/pSeAP-SBP. (**C**) Double knockout strains of E233Δ*cel2A*Δ*lacS*, E233Δ*cel1*Δ*lacS* and E233Δ*cel1*Δ*cel2A*. (**D**) Wild type REY15A and E233-pSeAP cultured in media with or without uracil (U). The RT-qPCR experiments were performed in triplicates, with TBP as the internal reference.

It was reported that cellodextrins (G2 to G8) are transported into cellular by ABC transporter substrate-binding protein SSO3053 in *Sa. solfataricus* (Lalithambika et al., 2012). By homology comparison, we identified its homolog in *Sa. islandicus* is SiRe_2190 and designated it as SisSBP. We constructed *sbp* knockout strain (Δ*sbp*) and overexpression strain (pSeAP-SBP) and performed CMC induction experiments. The results showed that in Δ*sbp*, *cel1*, *cel2A*, and *lacS* were downregulated to 18.97%, 48.60%, and 32.04%, respectively, of that in E233 (Figure 5A). Whereas in pSeAP-SBP strain (SBP protein was overexpressed, see Supplementary Fig.1A), the transcription levels of *cel1*, *cel2A*, and *lacS* were upregulated to 204.30%, 174.57%, and 124.25%, respectively, of that in control pSeAP (Figure 5B). These results suggest that in Δ*sbp*, the deletion of *sbp* impair the transport of oligosaccharides into the cell, resulting in reduced level or loss of cello-oligosaccharides and the downregulation of all three cellulase genes. Conversely, overexpression of *sbp* enhanced oligosaccharide transport into the cell, leading to the upregulation of *cel1*, *cel2A*, and *lacS*. To further verify the hypothesis that oligosaccharide accumulation induces cellulase transcriptional upregulation, we used overexpression strain of LacS and performed CMC induction experiments. As expected, the results showed that *cel1* and *cel2A* transcription levels were downregulated to 67.11% and 73.72%, respectively, of that in E233 carrying pSeAP. Since *lacS* overexpression promotes the degradation of intracellular oligosaccharides, intracellular oligosaccharide accumulation may be reduced (Figure 5B). These findings further support the hypothesis that intracellular levels of oligosaccharides can regulate the transcription of cellulase genes. It would be intriguing to investigate how the cells sense cellular cello-oligosaccharide level and regulate the endoglucanase activity.

In addition, we constructed three double mutants of the cellulase genes and analyzed the transcription of the remaining cellulase gene in CMC medium by RT-PCR. As show in Figure 5C, the transcription levels of *cel1* in Δ*cel2A*Δ*lacS*, *cel2A* in Δ*cel1*Δ*lacS*, and *lacS* in Δ*cel1*Δ*cel2A* reduced to 14.27%, 35.62%, and 32.53%, respectively, of those of the corresponding genes in E233. This result supports that there exists a interplay among the three cellulases.

### 3.6 The transcription of cellulase genes are upregulated in strains having uracil synthesis pathway

During transcriptional analysis of the cellulases, we observed that the E233 carrying pSeAP exhibited higher transcription levels of *cel1*, *cel2A*, and *lacS* (Figure 5D) than those in E233. To cultivate E233, uracil was added in the medium, while no uracil was supplemented in the medium for E233 carrying pSeAP, which has a uracil synthesis cassette *pyrEF*. To investigate the underlying cause of this phenomenon, we first examined whether the lower cellulase transcription is due to the addition of uracil. For this purpose, we used the wild-type strain *Sa. islandicus* REY15A, which possesses an intact *pyrEF* in genome, allowing autonomous uracil biosynthesis and enabling growth in both uracil-supplemented and uracil-free media (Deng et al., 2009). Transcription of the cellulases on CMC medium was examined using REY15A strain in both CMC-TVU and CMC-TV media. Interestingly, in CMC-TVU medium, similar to E233 carrying pSeAP, REY15A exhibited a significant upregulation of *cel1*, *cel2A*, and *lacS*, with transcription levels reaching 428.95%, 262.27%, and 228.40% of those in E233, respectively (Figure 5D). Thus, the influence of addition of uracil is ruled out. Taken together, these findings suggested that the transcription of cellulase genes in CMC-containing medium is upregulated in strains carrying the *pyrEF* cassette either in the genome or in the plasmid.

### 3.7 The growth of different strains in the presence of CMC is closely associated with transcription of the cellulase genes

We determined the growth curves of different strains cultured in medium with D-glucose or CMC as the carbon sources (Figure 6). For any single-knockout strains, Δ*cel1*, Δ*cel2A*, Δ*lacS*, and Δ*sbp*, no significant differences in growth were observed in D-glucose medium compared to E233. However, when cultured in CMC-containing medium, Δ*cel1*, Δ*cel2A*, and Δ*sbp* strains grew faster than E233 (Figure 6A). When cultured in CMC-containing medium, all three double-knockout strains Δ*cel2A*Δ*lacS*, Δ*cel1*Δ*lacS*, and Δ*cel1*Δ*cel2A* grew much quicker than E233 (Figure 6B). Interestingly, all the overexpressing strains pSeAP-Cel1, pSeAP-Cel2A, pSeAP-LacS, and E233/pSeAP-SBP as well as the control pSeAP exhibited pronounced growth lag in CMC, with pSeAP-Cel2A the most pronounced (Figure 6C). Similarly, REY15A which possesses *pyrEF* on the chromosome, also exhibited a comparable growth lag in CMC medium, and this lag was not attributable to the absence of uracil in the medium (Figure 6D). However, this significant growth lag is associated with the presence of the *pyrEF* gene and is only observed in CMC-containing medium, not in D-glucose conditions (Figure 6D).

**Fig. 6.**
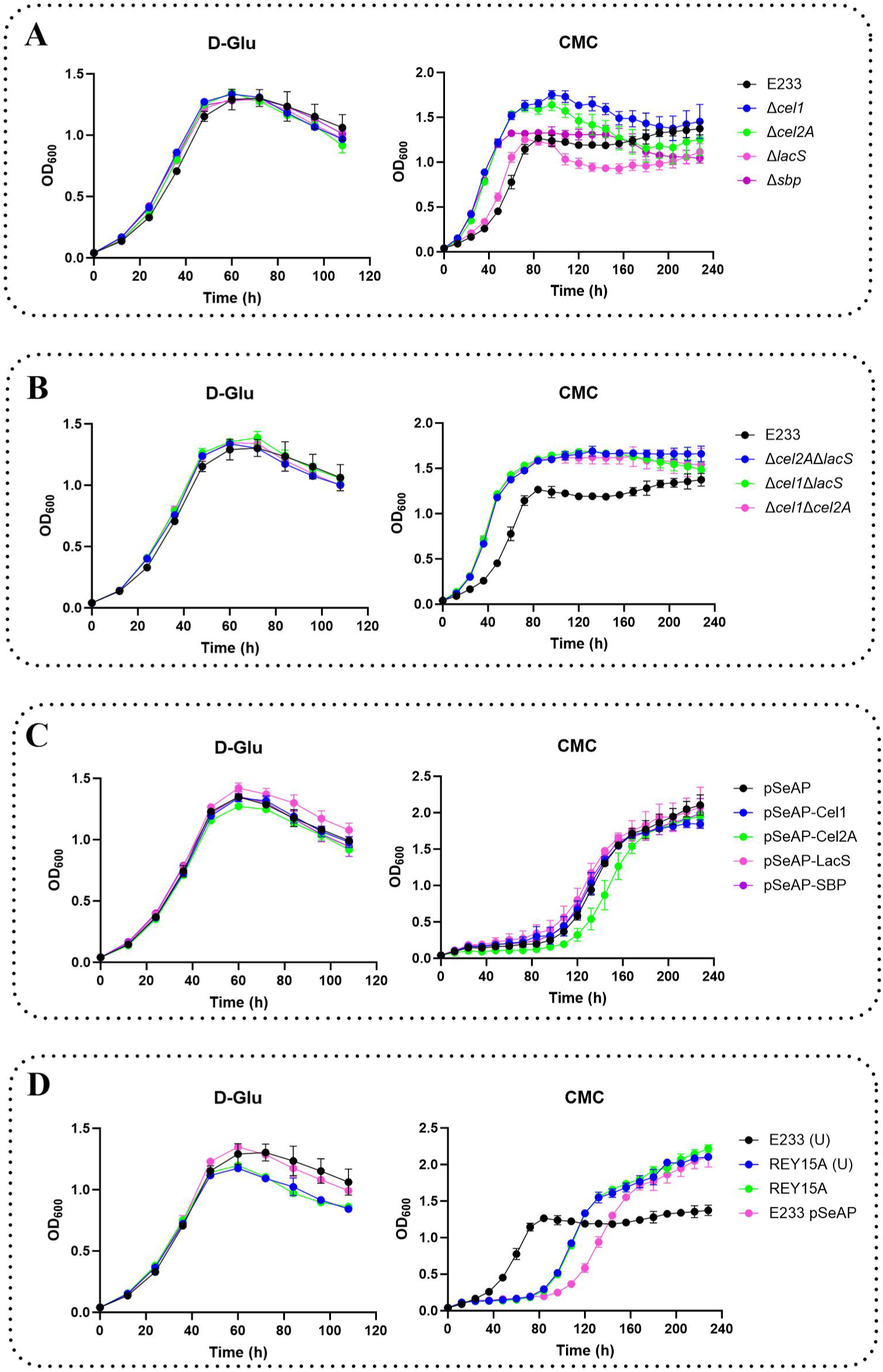
Growth curves of *Sa*. *islandicus* strains in different media. The cells were inoculated into 40 ml medium to a final estimated OD_600_ of 0.04 and the growth was monitored using spectrometer. (**A**) Sigle deletion mutants E233Δ*cel1*, E233Δ*cel2A*, E233Δ*lacS* and E233Δ*sbp* cultured in medium GTVU or CMC-TVU (TVU supplement with D-Glu or CMC). (**B**) Double deletion mutants E233Δ*cel2A*Δ*lacS*, E233Δ*cel1*Δ*lacS*, and E233Δ*cel1*Δ*cel2A* cultured in medium GTVU and CMC-TVU. (**C**) Overexpression strains of E233/pSeAP, E233/pSeAP-Cel1, E233/pSeAP-Cel2A, E233/pSeAP-LacS and E233/pSeAP-SBP cultured in medium GTV or CMC-TV (TV supplement with D-Glu or CMC). (**D**) REY15A and E233/pSeAP cultured in medium GTV(U) or CMC-TV(U).

By correlating the observed growth status with above cellulase transcription (Figure 5), we found that strains exhibiting downregulation of cellulase transcription, including Δ*cel1*, Δ*cel2A*, Δ*sbp*, Δ*cel2A*Δ*lacS*, Δ*cel1*Δ*lacS*, and Δ*cel1*Δ*cel2A*, consistently grew faster than those that exhibited higher cellulase transcription levels, including E233, REY15A, and strains carrying the plasmids (Figures 5 and 6). It appears that the downregulation of cellulase transcription is associated with cellular growth advantage, whereas the upregulation of cellulase transcription consistently coincides with growth lag. To further investigate the relationship between growth rate and cellulase transcriptional changes, we reduced the nutritional levels of the medium to 1/2, 1/4, and 1/8 of the original medium (Supplementary Figure 3A). The growth rates and the highest cell densities of E233 in the nutritional depleted medium were both slower than those in normal medium. As shown in Supplementary Figure 4, the transcription levels of the cellulase genes did not change compared with that in normal medium. This result suggests that growth rate variation alone was not the direct cause of significant cellulase transcriptional upregulation. Thus we conclude that overall upregulation of cellulases transcription leads to a growth retardance, but growth rate reduction does not directly result in cellulase upregulation.

## 4. Discussion

The microbes of *Sulfolobales* thrive in acidic and high-temperature environments, mostly sulfates containing hot springs, where nutrient input is limited (Brock et al., 1972). Nevertheless, many isolated strains are able to use 5 and 6-carbon sugar as carbon sources, which makes them as potential next generation of biomass degradation utilization platforms (Crosby et al., 2019; De Lise et al., 2023; Joshua et al., 2011). One of the standing questions is that the production and cellulase activity in the wild type cells are very low, and our initial experiments showed that it is difficult to express the cellulases (except the LacS), despite we have well-developed expression system in *Sa. islandicus* REY15A, highlighting that there exists an unknown mechanism for cellulase production, translocation, and regulation.

In this paper, we firstly constructed a vector pSeAP, which enables efficient and constitutive expression of LasS gene at high levels transcriptionally and translationally. We were able to detect the CMC activity and determine the localization of the endoglucanases (Figure 2 and 3). Our results confirm that both Cel1 and Cel2A are membrane-associated proteins, and the sizes of the two proteins detected were larger than their theoretical sizes (Figure 2B). This may be attributed to glycosylation modifications in the N-terminal transmembrane regions of these proteins, similar to what has been reported for the homologous protein SSO1354 of Cel2A in *Sa. solfataricus* (Girfoglio et al., 2012; Maurelli et al., 2008).

We have clarified that Cel2A is the only active endoglucanase in *Sa. islandicus* REY15A by gene knockout and gene expression. Our experiments demonstrated that Cel1 does not possess endoglucanase activity, which is inconsistent with the result of the homolog SSO2534 in *Sa. solfataricus* (Limauro et al., 2001), but in agreement with the study by Girfoglio et al. (Girfoglio et al., 2012), our result supports that the activity observed in colony of *Sa. solfataricus* is produced by SSO1354 and SSO1949 (Girfoglio et al., 2012; Huang et al., 2005). The result on the endoglucanase activity of Cel2A is consistent with that in *Sa. solfataricu* (Girfoglio et al., 2012; Huang et al., 2005).

Despite lack of endoglucanase activity for Cel1, we found that in CMC-containing medium, *cel1* was co-transcriptionally upregulated along with *cel2A* and *lacS* (Figure 4C) and deletion of *cell1* resulted in downregulation of the other two cellulase genes (Figure 5A and 5C), suggesting that Cel1 plays a role in CMC utilization. More generally, our gene knock out and over expression experiments showed that the transcription of the cellulases in *Sa. islandicus* are influenced by the deletion of one or two cellulases, or by the overexpression of *lacS* (Figure 5A-C). The results suggested that the three cellulases are interplayed. As for the mechanism, we assume that it would be dependent on intracellular oligosaccharides levels. Deletion of *lacS* and over expression of *sbp* make cellular intracellular oligosaccharide levels increase, this in turn stimulates the transcriptional upregulation of *cel1* and *cel2A*. Conversely, deletion of *sbp* or overexpression of *lacS* makes cellular intracellular oligosaccharide levels decrease, this in turn leads to transcriptional downregulation of *cel1* and *cell2A* (Figure 5A-B). It will be interesting to investigate how the cells sense the oligosaccharide level and regulate cellulase gene expression.

One of the interesting features in the cellulase transcriptional regulation is that *cel1* is regulated by aCcr1, a cell cycle regulator in *Sulfolobales* (Yang et al., 2023). This result suggests that cellulase gene transcription is coupled with cell cycle progression. Another accidental but interesting finding is that in the presence of *pyrEF*, transcription of all cellulase genes was upregulated (Figure 5D). The mechanism underlining this phenomenon is unclear. *pyrEF* is involved in the de novo synthesis pathway of uracil, and its intermediate products may affect intracellular sugar metabolism, leading to oligosaccharide accumulation. Alternatively, since uracil synthesis is essential for the cell in nature, its regulation should be controlled by cell cycle dependent transcription factors, in turn affecting cellulase gene transcription. Another intriguing finding is that there is a negative correlation between endoglucanase gene transcription and growth in CMC-containing medium (Figure 5 and 6). When *Sa. islandicus* was grown in CMC-containing medium, the cells exhibited significant growth advantages when cellulase transcription was downregulated. In contrast, when the cells harboring pSeAP or *pyrEF* cassette were grown under CMC-containing medium, *cell1* and *cell2A* transcription was enhanced, and a growth lag occurred (Figures 4C, 5D, and 6C and 6D). Collectively, our results suggest that there is complex and intricated mechanism for cellulose utilization in Sulfolobales archaea.

Based on these findings, we propose a model for cellulose utilization in *Sa islandicus* (Figure 7). Upon biomass is available, the cellulase genes are induced by oligosaccharides degraded by cellulases. The endoglucanase Cel2A, either in its membrane-anchored or secreted form, degrades extracellular long-chain cellulose into oligosaccharides, which are then transported into the cell via the ABC transporter system and subsequently hydrolyzed into monosaccharides by intracellular LacS. The transcriptional regulator aCcr1 directly regulates the transcription of *cel1*. Additionally, the transcription of *cel1*, *cel2A*, and *lacS* are influenced by intracellular oligosaccharide levels and interplay among the cellulases. Furthermore, uracil synthesis by *pyrEF* enhances cellulase gene transcription by an unknown mechanism. Once biomass is depleted, cellular oligosaccharide level decreases, leading to reduced cellulase transcription to save energy for cell propagation. By this ingenious and intricated mechanism, the cells are adapted to the nutrient-limited extreme environments.

**Fig. 7.**
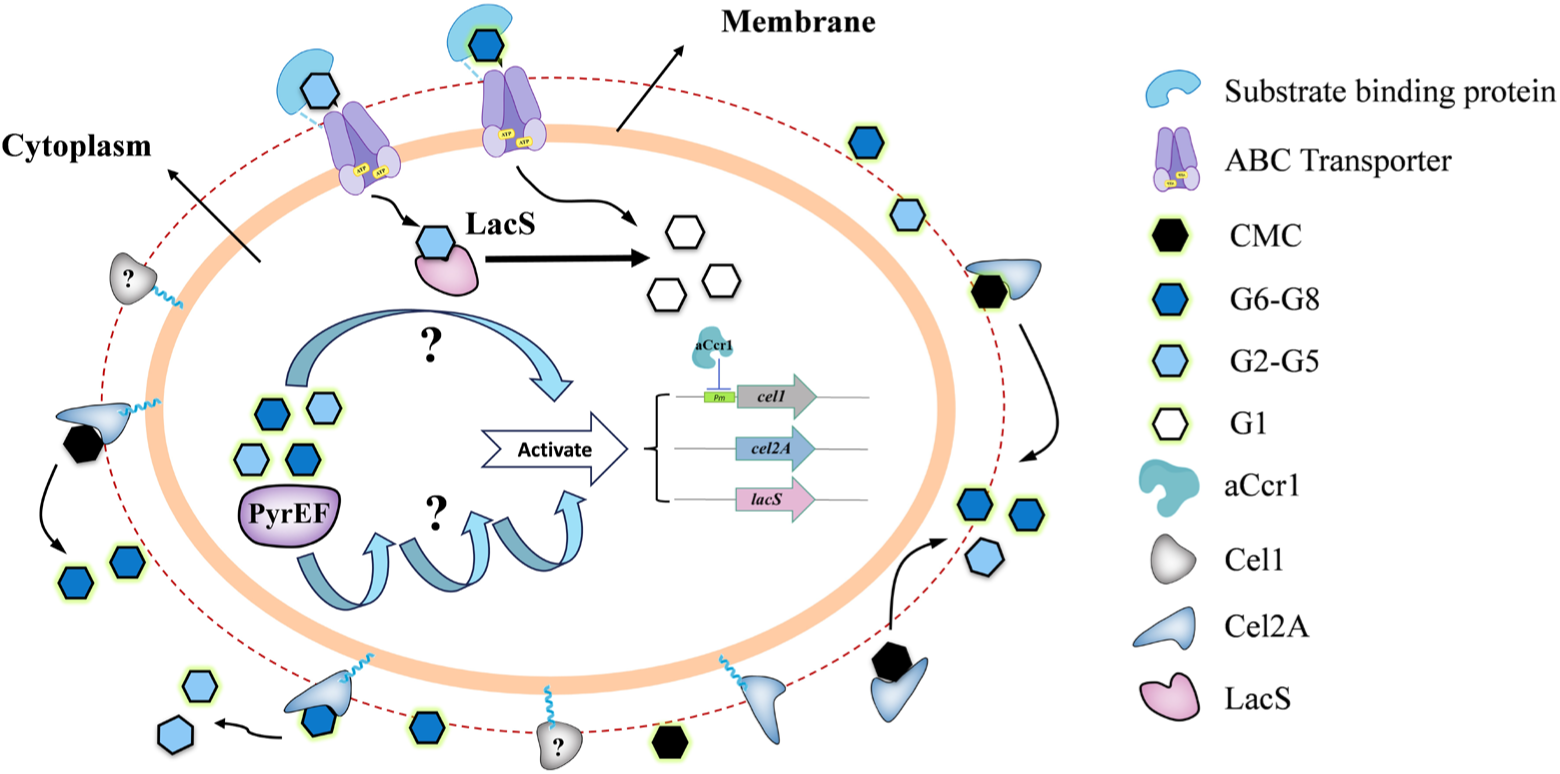
A model for cellulose utilization in Sulfolobales archaea. Upon biomass is available, the cellulase genes are induced by oligosaccharides degraded by cellulases. The endoglucanase Cel2A, either in its membrane-anchored or secreted form, degrades extracellular long-chain CMC into oligosaccharides, which are then transported into the cell via the ABC transporter system and subsequently hydrolyzed into monosaccharides by intracellular LacS. The transcriptional regulator aCcr1 directly regulates the transcription of *cel1*. Additionally, the transcription of *cel1*, *cel2A*, and *lacS* are influenced by intracellular oligosaccharide levels. Furthermore, uracil synthesis by *pyrEF* enhances cellulase gene transcription by an unknown mechanism.

## 5. Conclusion

In this study, we constructed a constitutive expression vector pSeAP for *Sa. saccharolobus*. Detailed investigation was carried out on three cellulases in *Sa. islandicus*. It was revealed that both Cel1 and Cel2A are membrane proteins, and Cel2A is the only active endoglucanase in *Sa. islandicus*. The transcription of cellulases is regulated by many factors, including the transcription factor aCcr1, CMC, and *pyrEF*. The transcription of cellulases in S*a. islandicus* is probably affected by intracellular oligosaccharide levels and there may exist connections between the transcription of cellulases and the growth status in CMC-containing medium. A transcriptional induction model of cellulases in *Sa. islandicus* is proposed. This work provides new insights into the roles and mechanism of cellulose degradation by cellulases in acidothermophilic archaea and lays a foundation for engineering Sulfolobales cells for synthetic biology.

## Notes

### Competing Interest Statement

The authors have declared no competing interest.

